# An inhibitory acetylcholine receptor gates context dependent mechanosensory processing in *C. elegans*

**DOI:** 10.1101/2024.03.21.586204

**Authors:** Sandeep Kumar, Anuj K. Sharma, Andrew M. Leifer

## Abstract

An animal’s current behavior influences its response to sensory stimuli, but the molecular and circuit-level mechanisms of this context-dependent decision-making is not well understood. In the nematode *C. elegans*, inhibitory feedback from turning associated neurons alter downstream mechanosensory processing to gate the animal’s response to stimuli depending on whether the animal is turning or moving forward [1–3]. Until now, the specific neurons and receptors that mediate this inhibitory feedback were not known. We use genetic manipulations, single-cell rescue experiments and high-throughput closed-loop optogenetic perturbations during behavior to reveal the specific neuron and receptor responsible for receiving inhibition and altering sensorimotor processing. An inhibitory acetylcholine gated chloride channel comprised of *lgc-47* and *acc-1* expressed in neuron RIM receives inhibitory signals from turning neurons and performs the gating that disrupts the worm’s mechanosensory evoked reversal response.

---

The nematode *C. elegans*’s response to mechanosensory stimuli is influenced by its current behavior: it is less likely to reverse in response to a stimulus that it receives when in the midst of a turn compared to a stimulus that it receives when moving forward Fig 1A [2]. Because turns are fast, typically lasting approximately two seconds, we hypothesize that synaptic signaling via neurons may mediate this context dependency. Understanding the molecular and circuit mechanism of this sensorimotor processing will provide insights into how a simple nervous system implements context-dependent decision-making.

**Figure 1.**
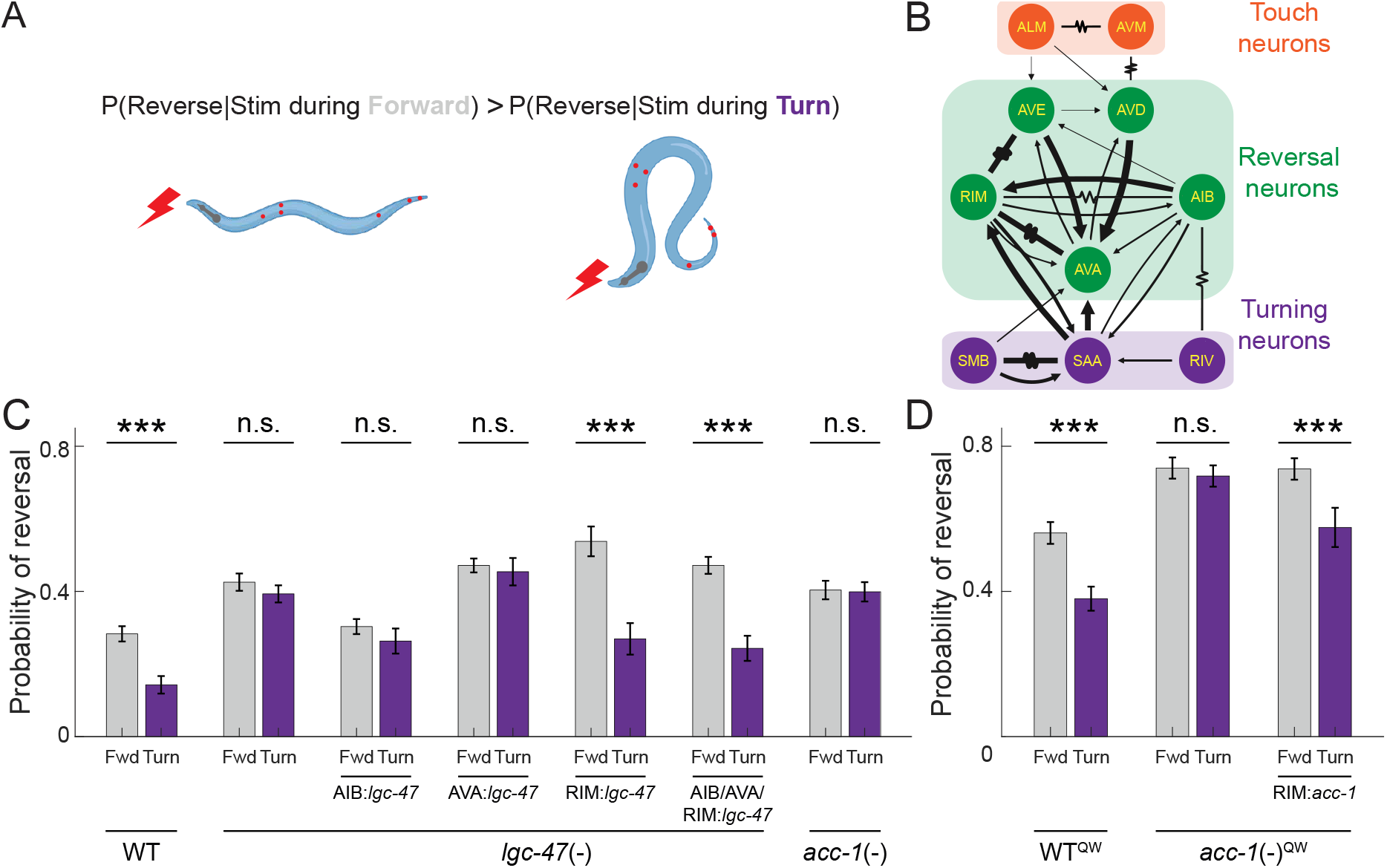
LGC-47 and ACC-1 in neuron RIM gate turning-dependent reversals. A) Optogenetic stimulation is delivered to a worm expressing Chrimson in the six gentle touch neurons during forward or turning locomotion. B) Anatomical wiring diagram of touch receptor neurons, turning-associated neurons and reversal-associated interneurons (adapted from Nemanode [11]). C) Probability of reversal in response to optogenetic stimulation delivered during forward movement or turning onset is shown for different genetic manipulations, including loss-of-function of *lgc-47* or *acc-1*, and their corresponding cell-specific rescues. All animals express Chrimson under *mec-4* promoter including “WT.” N > 400 stim events for each condition (Fig S1). D) Reversal response for *acc-1* loss-of-function mutants and corresponding cell-specific rescue. Here a different *mec-4* Chrimson allele is used, denoted by ^QW^. N > 320 stimulation events per condition (Fig S1). ^***^ indicates p < 0.001, ‘n.s.’ indicates p > 0.05 via two proportion Z-test.

The gentle touch mechanosensory circuit in *C. elegans* has been extensively studied [4] and its downstream interneurons are known Fig. 1B. When the worm turns, inhibitory feed-back from turning neurons is thought to disrupt downstream processing of mechanosensory signals by inhibiting some of these interneurons [3]. In particular, inhibitory feedback from a collection of turning-associated neurons SMB, SAA and RIV [5], decrease the likelihood of reversing during a turn [3], but the exact location and mechanism with which these turning signals interact with downstream mechanosensory processing is unknown. Previously we performed optogenetic activation studies showing that inhibitory signals from turning neurons arrive at or upstream of reversal interneuron AVA, because only activation of those neurons, and not AVA evoked reversals in a turning-dependent manner [3]. Here we seek to identify the neurons and receptors that receive inhibition from the turning circuit.

We investigated the outgoing synapses of SAA because that neuron type is known to be involved in turning [3, 5, 6] and it makes synaptic contacts onto reversal neurons including AVA, RIM and AIB [7, 8], Fig 1B. We specifically investigated the role of inhibitory acetylcholine receptors expressed at these synapses because SAA releases acetylcholine [9–11] and we expect it to send an inhibitory signal [3, 5]. We focused on *lgc-47* and *acc-1*, two genes for inhibitory acetylcholine receptors, with known expression in neurons RIM, AIB, and AVA among others [10, 11]. We investigated LGC-47 first because it expresses at higher levels than ACC-1, Supplementary Figure S1A [11]. We therefore measured the response to mechanosensory stimuli in *lgc-47* loss-of-function mutants.

We expressed the light-gated ion channel Chrimson in the six gentle touch receptor neurons under the control of a *mec-4* promoter, and used a high throughput closed-loop optogenetic delivery system to automatically stimulate animals either when moving forward or triggered upon the onset of a turn [2], Fig 1C. *lgc-47* loss-of-function mutants exhibited little or no turning-dependent gating of the reversal response: they were similarly likely to reverse in response to stimuli regardless of whether the stimulus was delivered while the animal was turning or moving forward. By contrast, wild-type background animals showed significant gating: they were significantly less likely to reverse in response to stimuli delivered while turning compared to stimuli delivered during forward movement. These measurements suggest that LGC-47 mediates gating of mechanosensory evoked reversals.

To identify where LGC-47 acts, we performed cell-specific rescues in the reversal associated interneurons AIB, AVA, RIM or all three, by expressing WT *lgc-47* cDNA in the background of *lgc-47* loss-of-function animals, Supplementary Table S1. For each rescue, we measured the animal’s response to optogenetically induced mechanosensory stimuli, Fig 1C. Only animals that expressed LGC-47 in RIM recapitulated the WT gating behavior. We therefore conclude that LGC-47 mediates the gating of mechanosensory evoked reversals by inhibiting reversal neuron RIM in response to acetylcholine release from SAA. In other words, SAA inhibits RIM via LGC-47 during turns to prevent the initiation of a reversal.

We next investigated the candidate inhibitory acetylcholine receptor gene *acc-1* that is expressed at lower levels in a similar pattern of neurons [9, 12]. Intriguingly, recent work suggests that ACC-1 regulates the duration of spontaneous reversals in a manner similar to what we proposed for LGC-47: namely ACC-1 is thought to inhibit RIM upon SAA activation, in that case halting an ongoing reversal [13]. We wondered whether ACC-1 also contributes to turning-dependent gating by potentially stopping reversals before they start.

*acc-1* loss-of-function mutants showed no turning-dependence in their mechanosensory evoked responses, Fig 1 C,D, just like *lgc-47* mutants. We observed the same effect both in our *mec-4::Chrimson* background strains Fig 1 C and in a nominally similar but separately generated set of strains from the Quan Wen group, Fig 1 D. Turning dependence reappeared when ACC-1 was rescued in the neuron type RIM, just as it did for LGC-47. Therefore ACC-1 performs the same role as LGC-47 in mediating the turning dependent gating. Recent *in vitro* electrophysiology studies suggest that LGC-47 and ACC-1 may form a heteromeric ion channel [14], and our finding that LGC-47 and ACC-1 perform the same role *in vivo* supports this hypothesis.

Taken together, we conclude that neuron type SAA gates mechanosensory evoked reversals by inhibiting reversal neurons RIM via an inhibitory acetylcholine receptor comprised of LGC-47 and ACC-1. Inhibition of RIM during turns is ideally situated to prevent reversals because it makes many gap junctions with neurons AVA and AVE, as well as gap junctions with AIB— all neurons implicated in promoting reversals. Inhibition of RIM may serve as a shunt to inhibit activity across the reversal circuitry. Future imaging studies are needed to reveal the neural dynamics of these neurons in response to mechanosensory stimuli delivered during turns.

## DIVERSITY AND INCLUSION STATEMENT

We support inclusive, diverse, and equitable conduct of research.

## Supplemental Text

### Strains

All strains used in this work are listed in Table S1. The CRISPR engineered null mutant strain PS8742 [*lgc-47(sy1501)*] for the LGC-47 receptor was obtained from the Caenorhabditis Genetics Center (CGC). The mutant strain CX12721 [*acc-1(tm3268)*] defective for the ACC-1 receptor was a gift from Dr. Cori Bargmann, Rockefeller University. To rescue the *lgc-47* cDNA in neurons RIM, AIB, and AVA, we used the cell specific promoters *tdc-1P, npr-9P*, and *rig-3P* respectively. Strains WEN0920, WEN1015, and WEN1025, were gifts from Dr. Quan Wen, University of Science and Technology of China. RNA expression levels of *lgc-47* and *acc-1* were reported from the CeNGEN database [11] and displayed in SFig S1A.

### Nematode handling

Worm handling was performed as described in [3]. Briefly, all the strains used in this study were grown on standard nematode growth media plates with OP-50 (*E. coli*) as a food source at 20 C. Agarose plates containing gravid worms were bleached to collect eggs. The eggs were rinsed with M9 solution at least three times and left on a shaker overnight. The next day, L1 larvae were plated on a freshly seeded plate containing OP50 mixed with 1 ml of 0.5 mM all-trans-retinal and placed in a dark container in a 20 C incubator until day 1 young adult stage, at which time experiments were performed.

### Optogenetic stimulation

To measure the response of mechanosensory stimulus during forward and turning behaviors, we delivered optogenetic stimulus to the worms using a high throughput optogenetic delivery system [2]. Optogenetic stimulation was performed as described previously [3]. Briefly, an open-loop optogenetic stimulation protocol was used to stimulate the animal when the animal was moving forward. 3 s of 80 uW/mm^2^ of 630 nm illumination was delivered to all animals on the plate every 30 s. Only stimuli that landed when the animal was moving forward was considered. To investigate the behavioral response to stimuli during turns, a closed-loop behavior-triggered stimulus was used. 3 s of illumination was delivered to an animal whenever the system detected that a worm was initiating a turn, but no more often than once every 30 s to the same animal.

### Behavior Analysis

Behavior classification of forward movements, turns and reversals was performed as described previously [3]. Briefly, two sets of behavior mapping algorithms were used in this study, one for real-time tracking of worms and optogenetic stimulation, and another more conservative one for post-processing analysis. The real-time algorithm tracked each worm as it was crawling on agarose plates and determined locomotory parameters in real-time such as velocity, centerline, body curvature, etc. When the system detects that the worm is initiating a turn, a computer controlled projector delivers an optogenetic stimulus precisely to that worm. In post-processing, we classify the worm’s behavior, and determine whether the worm reversed in response to stimuli [1, 15]. We exclude worms that do not move for prolonged periods of time. A worm is classified as reversing in response to stimuli if its velocity is less than or equal to -0.11 mm/s during the 3 second optogenetic stimulation window. Details of the number of experimental assays and stimulus events can be found in Fig S1B.

## Statistical Analysis

In our analysis, stimulus events are the fundamental units, and we calculate statistics based on the probability of exhibiting a response to a stimulus event. We report three statistics: the proportion of stimulus events that resulted in a reversal of the worm, the total number of stimulus events presented to the worm, and the corresponding 95% confidence interval for the proportions of the worm reversing. To reject the null hypothesis that two empirically observed proportions (for example during forward and turning) are the same, we used a two-proportion Z-test and reported a p-value. A p-value<0.05 was considered significant.

## Data and code availability

Computer-readable files showing processed tracked behavior and optogenetic stimulus events for all experiments are publicly available at https://doi.org/10.6084/m9.figshare.25396453. All analysis codes used in this manuscript are available at https://github.com/leiferlab/kumar-molecular-mechanism.git.

## Strains and plasmid availability

Transgenic strains AML67, AML597, AML614, AML617, AML618, AML622, and AML627 will be made publicly available through the Caenorhabditis Genetics Center (CGC).

## Acknowledgements

We thank Quan Wen (University of Science and Technology of China) for providing us the genetic strains: QW0920, QW1015, and QW1025. We also thank Dr. Cori Bargmann for providing us the CX12721 strain. This work used computing resources from the Princeton Institute for Computational Science and Engineering. The strains of this work are distributed by the CGC, which is funded by the NIH Office of Research Infrastructure Programs (P40 OD010440). The research reported in this work was supported by the National Science Foundation (https://www.nsf.gov) through an NSF CAREER Award to AML (IOS-1845137) and through the Center for the Physics of Biological Function (PHY-1734030); and by the National Institute of Neurological Disorders and Stroke (https://www.ninds.nih.gov/) of the National Institutes of Health, National Institute of Neurological Disorder and Stroke under New Innovator award number DP2-NS116768 to AML; and by the Simons Foundation (https://www.simonsfoundation.org/) under award SCGB 543003 to AML. The funders had no role in study design, data collection and analysis, decision to publish, or preparation of the manuscript. The content is solely the responsibility of the authors and does not represent the official views of any funding agency.

## Author Contributions

Conceptualization: A.M.L. and S.K.; Formal analysis: S.K.; Funding acquisition: A.M.L.; Investigation: S.K.; Methodology: S.K. and A.K.S.; Project administration: A.M.L.; Resources: A.K.S.; Supervision: A.M.L.; Writing – original draft: S.K.; Writing - review & editing: A.M.L.,S.K and A.K.S.

## Competing Interests

The authors declare no competing interests.

## Supplemental Figures

**Figure S1.**
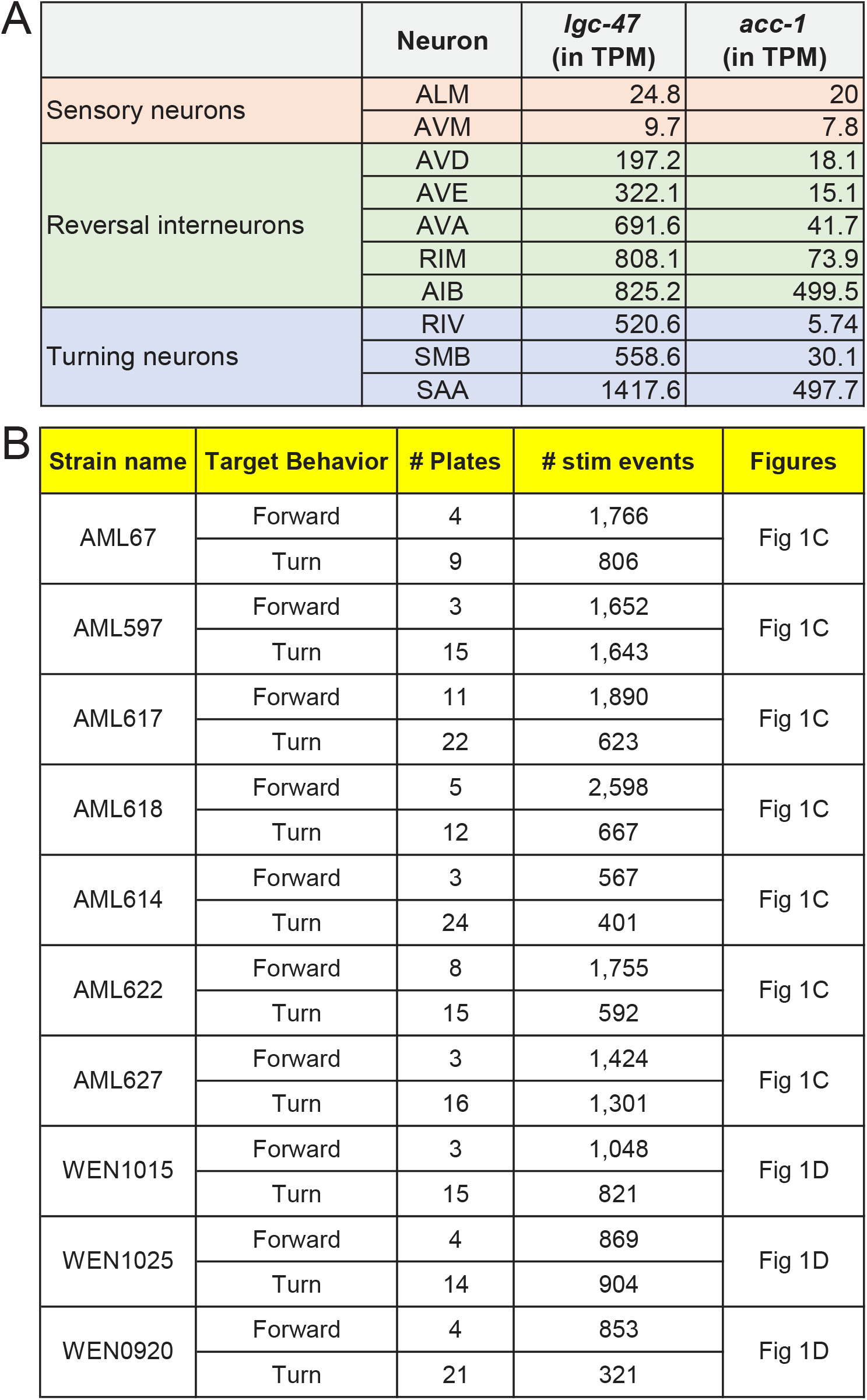
Reported receptor expression levels and details of experiments performed. A) Relative expression levels of *lgc-47* and *acc-1* in transcripts per million (TPM) as reported in [CeNGEN](https://cengen.shinyapps.io/CengenApp/) single cell gene expression atlas [11, 16] (no threshold was selected). B) Strains used, number of plates and number of stimulation events are listed for each experiment.

## Supplemental Tables

**Table S1.** Strains used. Genotype of strains used in this study.

| Strain name | Expression | Genotype | Background | Figure | Ref |
| --- | --- | --- | --- | --- | --- |
| AML67 | Chrimson in ALML/R, AVM, PLML/R, PVM | *wtfIs46[Pmec-4::Chrimson::SL2::mCherry::unc-54 40ng/ul]* | N2-WT | Fig 1C | [1] |
| AML597 | Chrimson in ALML/R, AVM, PLML/R, PVM | *lgc-47(sy1501) X; wtfIs46[mec-4P::Chrimson::SL2::mCherry::unc-54 40 ng/ul]* | *lgc-47 (sy1501)* | Fig 1C | This work |
| AML617 | Chrimson in ALML/R, AVM, PLML/R, PVM; rescuing *lgc-47* in AIB neuron | *lgc-47(sy1501) X; wtfIs46[mec-4P::Chrimson::SL2::mCherry::unc-54 40 ng/ul]; wtfEX538 [npr-9P::Al::lgc-47::SL2::tagBFP 30ng/ul + Coel::GFP 70ng/ul]* | *lgc-47 (sy1501)* | Fig 1C | This work |
| AML618 | Chrimson in ALML/R, AVM, PLML/R, PVM; rescuing *lgc-47* in AVA neuron | *lgc-47(sy1501) X; wtfIs46[mec-4P::Chrimson::SL2::mCherry::unc-54 40 ng/ul]; wtfEX539 [rig-3P::Al::lgc-47::SL2::GFP 30ng/ul + Coel::GFP 70ng/ul]* | *lgc-47 (sy1501)* | Fig 1C | This work |
| AML614 | Chrimson in ALML/R, AVM, PLML/R, PVM; rescuing *lgc-47* in RIM neuron | *lgc-47(sy1501) X; wtfIs46[mec-4P::Chrimson::SL2::mCherry::unc-54 40 ng/ul]; wtfEX535 [tdc-1P::Al::lgc-47::SL2::his-24::tagRFP 30ng/ul + Coel::GFP 70ng/ul]* | *lgc-47 (sy1501)* | Fig 1C | This work |
| AML622 | Chrimson in ALML/R, AVM, PLML/R, PVM; rescuing *lgc-47* in AIB, AVA, and RIM neuron | *lgc-47(sy1501) X; wtfIs46[mec-4P::Chrimson::SL2::mCherry::unc-54 40 ng/ul]; wtfEX543 [tdc-1P::Al::lgc-47::SL2::his-24::tagRFP 30ng/ul + npr-9P::Al::lgc-47::SL2::tagBFP 30ng/ul + rig-3P::Al::lgc-47::SL2::GFP 30ng/ul + Coel::GFP 70ng/ul]* | *lgc-47 (sy1501)* | Fig 1C | This work |
| AML627 | Chrimson in ALML/R, AVM, PLML/R, PVM | *acc-1 (tm3268)IV; wtfIs46[mec-4P::Chrimson::SL2::mCherry::unc-54 40 ng/ul]* | *acc-1 (tm3268)* | Fig 1C | This work |
| WEN1015 | Chrimson in ALML/R, AVM, PLML/R, PVM | *wenIs1015[Pmec-4::chrimson::mcherry(quan0047,50ng/ul), Plin-44::gfp]* | N2-WT | Fig 1D | [13] |
| WEN1025 | Chrimson in ALML/R, AVM, PLML/R, PVM | *acc-1(tm3268); wenIs1015[Pmec-4::Chrimson::mCherry; Plin-44::GFP]* | *acc-1 (tm3268)* | Fig 1D | [13] |
| WEN0920 | Chrimson in ALML/R, AVM, PLML/R, PVM; rescuing *acc-1* in RIM | *acc-1(tm3268); wenIs1015[Pmec-4::Chrimson::mCherry; Plin-44::GFP]; wenEx0920[Ptdc1::acc1::GFP(20ng/ul); Plin-44::mcherry]* | *acc-1 (tm3268)* | Fig 1D | [13] |

## Notes

### Competing Interest Statement

The authors have declared no competing interest.

